# Cells Engage Endogenous Malonate Synthesis to Drive Mitochondrial Metabolism

**DOI:** 10.64898/2026.05.22.727248

**Authors:** Riley J. Wedan, Pieter R. Norden, Morgan T. Canfield, Abigail E. Ellis, Sanskriti Saxena, Jacob Z. Longenecker, Michelle Dykstra, Ryan D. Sheldon, Sara M. Nowinski

## Abstract

Malonate is often described as an endogenous inhibitor of complex II of the electron transport chain. However, the cellular source of malonate is unclear, and current knowledge concerning its metabolism is limited to the action of a single enzyme, Acyl-CoA Synthetase Family Member 3 (ACSF3), which converts malonate to malonyl-CoA in the mitochondrial matrix. One potential route of malonate metabolism downstream of ACSF3 is its consumption by the mitochondrial fatty acid synthesis (mtFAS) pathway. However, studies examining the link between ACSF3 and mtFAS have yielded conflicting results. We developed a novel mass spectrometry approach to perform stable isotope tracing into products of mtFAS, and found that while malonate is in fact a carbon source for mtFAS, ACSF3 is not required for malonate incorporation into mtFAS products. Using this method to trace other nutrients into mtFAS, we also found evidence of acetyl-CoA carboxylase 1 (ACC1)-dependent malonate synthesis from glucose. We further show that ACC1 is required for optimal mtFAS activity, with downstream effects on oxidative phosphorylation. Together these findings establish the malonate as a regulated endogenous intermediate that supports mtFAS activity and mitochondrial oxidative function.

## INTRODUCTION

Malonate has recently gained attention for its potential use as a treatment for myocardial infarction to prevent ischemia-reperfusion injury (IRI)^1,2^. In IRI, unregulated succinate oxidation results in catastrophic reactive oxygen species (ROS) production by the mitochondrial electron transport chain (ETC). Malonate is a competitive inhibitor of succinate dehydrogenase, also known as complex II of the ETC, and thus is thought to reduce the burden of ROS during reperfusion^2^. However, complex II and the ETC as a whole play other important roles in ATP production, the maintenance of redox balance, and the regulation of apoptosis^3^. Therefore, malonate is classically considered to be a toxic endogenous metabolite. In fact, malonate accumulates in metabolic disorders such as combined malonic and methylmalonic aciduria (CMAMMA), where it causes damage to tissues through an unknown mechanism^4^. Thus, as malonate begins to be more widely explored as a therapeutic, important considerations will need to be given to its endogenous role(s) and regulation.

Although the toxic effects of malonate accumulation have long been documented, the source of endogenous malonate is unclear. Dietary and microbiome sources, decarboxylation of oxaloacetate, or removal of coenzyme A (CoA) from malonyl-CoA have all been proposed, but the exact origins and metabolic pathways that create cellular malonate have remained elusive^5^. Moreover, how malonate is metabolized is similarly poorly understood. The only known mechanism for malonate detoxification in mammals is through the action of Acyl-CoA Synthetase Family Member 3 (ACSF3), the enzyme that is dysfunctional in CMAMMA. ACSF3 converts malonate to malonyl-CoA in the mitochondrial matrix, where it has been associated with protein malonylation^5^. While protein malonylation has been shown to regulate protein structure and function^6^, it is unclear the extent to which this pathway can adapt to detoxify large doses of exogenous malonate, and what other pathways for malonate metabolism might exist.

Unlike malonate, malonyl-CoA cannot cross mitochondrial membranes, and therefore once produced in the mitochondrial matrix by ACSF3, malonyl-CoA must be further metabolized within the mitochondria. Classically, malonyl-CoA levels in the cytoplasm are tightly regulated by acetyl-CoA carboxylases (ACC1 and ACC2), and malonyl-CoA decarboxylase (MLYCD), to control the rate of cytoplasmic fatty acid synthesis(FAS)^7^. In parallel to cytoplasmic FAS, *de novo* synthesis of fatty acids occurs in mitochondria through the mitochondrial fatty acid synthesis (mtFAS) pathway. Like cytoplasmic FAS, mtFAS requires malonyl-CoA as a carbon source^8^, and thus mtFAS is a likely endpoint for malonate metabolized by ACSF3. However, there is some confusion in the literature as to whether ACSF3 is required for mtFAS, with some describing it as the first step in the pathway while others find no effect of ACSF3 loss on mtFAS-dependent endpoints^9,10^.

To understand the metabolic origin of malonate, its metabolism, and its functional role in mammalian cells, we developed a novel carbon-tracing approach to directly assess carbon contributions to mtFAS pathway products by mass spectrometry. Using this approach, we found that exogenous malonate is readily incorporated into mtFAS-derived acyl chains, demonstrating that malonate can serve as a carbon source for this pathway. However, ACSF3 expression was not required for malonate incorporation into mtFAS products. We further found that when exogenous malonate is absent, or present at low, physiologic levels, cells instead rewire metabolism to sustain mtFAS activity from glucose. Unexpectedly, we discovered that cells turn on synthesis of endogenous malonate from glucose in an ACC1-dependent manner to support this pathway in the absence of supraphysiologic malonate. Together, our findings establish malonate as a regulated metabolic intermediate rather than a simple toxic byproduct, define its role in supporting mtFAS activity, and challenge the prevailing model that ACSF3-mediated malonyl-CoA production is the dominant route connecting malonate to the mtFAS pathway.

## RESULTS

### Malonate drives mitochondrial fatty acid synthesis

The only described fate of malonate intracellularly is its conversion to malonyl-CoA by the mitochondrial enzyme ACSF3^5,11^. ACSF3-derived mitochondrial malonyl-CoA has long been proposed to be the source of substrate for mitochondrial fatty acid synthesis (mtFAS) due to compartmental overlap^11^. However, this study and others have noted no effect of ACSF3 knockdown or knockout on mtFAS-dependent endpoints, raising questions about whether this is indeed the case^9,11^. mtFAS is a Type II fatty acid synthesis system that builds fatty acyl chains on the mitochondrial acyl carrier protein, NDUFAB1. mtFAS-derived acyl chains directly support iron sulfur cluster biogenesis and electron transport chain (ETC) assembly, or are further modified to form lipoic acid, a required co-factor for TCA cycle enzyme function and amino acid catabolism^12^. Activity of the mtFAS pathway has most often been monitored by western blot, probing for protein lipoylation and SDHB stability, two indirect downstream endpoints that depend on synthesis of mtFAS products. Indeed, deficiencies in *Mcat*, the first gene in the mtFAS pathway, result in loss of both endpoints in both mouse myoblasts (C2C12, Figure 1A) and rat cardiac myoblasts (H9c2, Figure 1B, S1A). SDHB stability is a surrogate for a larger ETC assembly defect, which is evident by blue native PAGE (Figure 1C-D and S1B). These phenotypes are all rescued by re-expression of Mcat.

**Figure 1.**
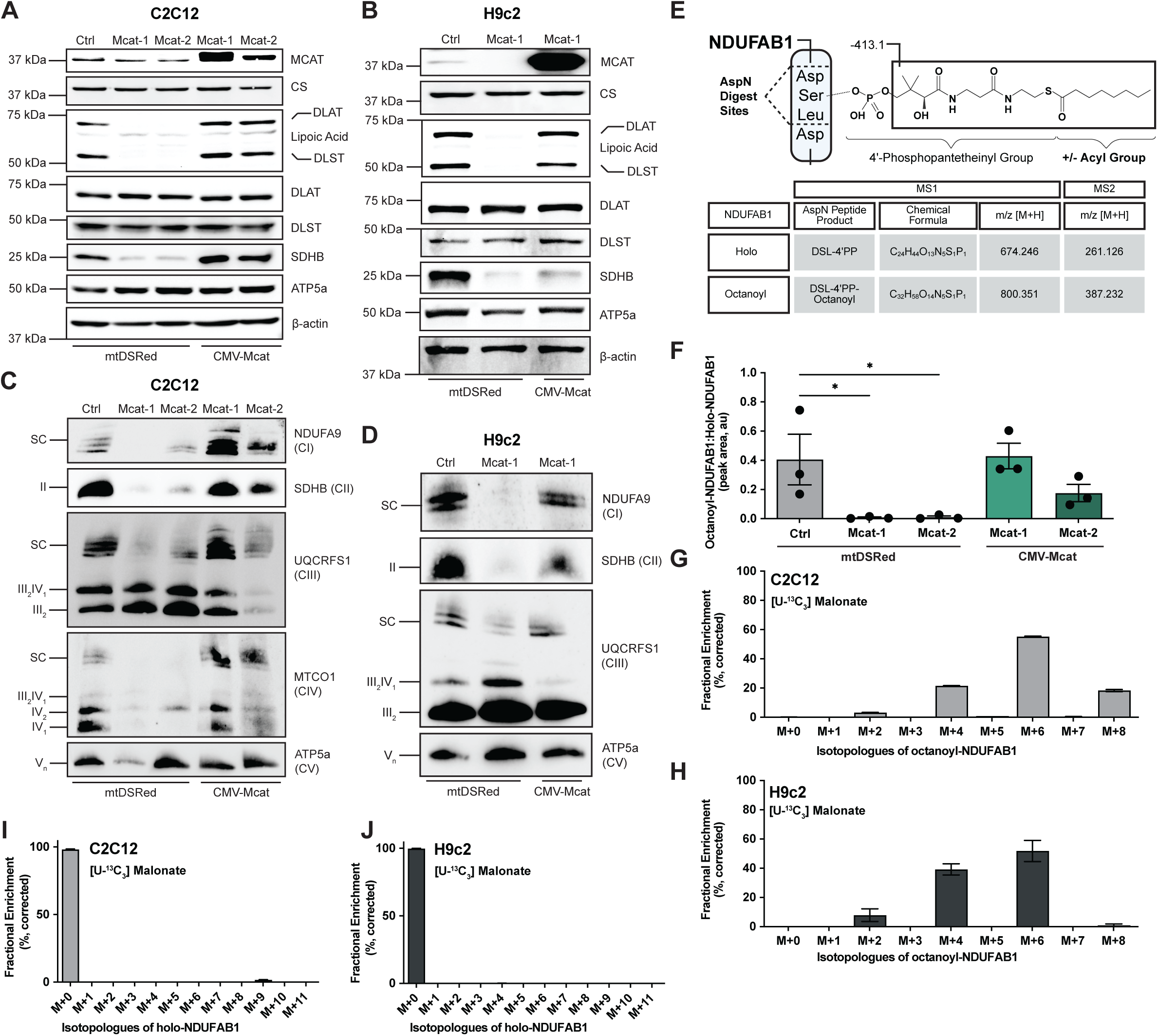
Malonate Drives Mitochondrial Fatty Acid Synthesis. **(A-B)** Whole cell lysates or mitochondrial enriched lysates (MCAT, CS blots) from (A) C2C12 mouse myoblast or (B) H9c2 rat cardiac myoblast clonal cell lines of the indicated genotypes expressing mtDSRed or Mcat were immunoblotted for the indicated protein targets. Data are representative of 3 biological replicates. Ctrl = control, CS = citrate synthase, DLAT = dihydrolipoamide s-acetyltransferase, DLST = dihydrolipoamide S-succinyltransferase, SDHB = succinate dehydrogenase B, ATP5a = mitochondrial loading control, β actin = whole cell loading control (**C-D**) Blue-Native PAGE of crude isolated mitochondrial lysates from (C) C2C12 and (D) H9c2 clonal cell lines of the indicated genotypes expressing mtDSRed or Mcat, immunoblotted for the indicated ETC complex subunits. Data are representative of 3 biological replicates. (**E**) Diagram depicting peptide products produced by digest of NDUFAB1 with AspN endoproteinase and resulting ions detected by mass spectrometry. (**F**) Relative abundance of octanoyl-NDUFAB1 normalized to holo-NDUFAB1 in the indicated cell lines. Statistical analysis was performed by one-way ANOVA followed by Tukey’s post-hoc test, * = p<0.05 (n=3). (**G-H**) Mass isotopologue distribution of octanoyl-NDUFAB1 in (G) wild-type C2C12 cells or (H) wild-type H9c2 cells after culture with uniformly labeled U-^13^C_3_ malonate for 3 hours (n=3). (**I-J**) Mass isotopologue distribution of holo-NDUFAB1 in (I) wild-type C2C12 cells or (J) wild-type H9c2 cells after culture with uniformly labeled malonate for 3 hours (n=3). Error bars represent +/- 1 SEM.

While convenient, these blotting-based assays are indirect and rely on turnover of covalent modifications, rendering real-time assessment of changes in mtFAS pathway activity impossible. Thus, to directly test mtFAS activity we turned to detecting modifications on NDUFAB1 via mass spectrometry^13–15^ (Figure 1E). FLAG-tagged NDUFAB1 was expressed in the cell lines of interest (Figure S1C-E), immunoprecipitated, and digested with the endoproteinase AspN, generating a short peptide product that spans the acyl site attachment as previously described^14^. This digest product was then detected by unique mass identifiers for acylated (octanoyl, 8:0) compared to holo-NDUFAB1. We detected identical peaks in digested, immunoprecipitated NDUFAB1-FLAG and digested standards, with no signal observed in digested immunoprecipitates from cells lacking NDUFAB1-FLAG (Figure S1F). To increase sensitivity and specificity of detection and enable carbon tracing experiments using stable isotope labeling, we used parallel-reaction monitoring (PRM) with a wide isolation window conducive to detecting all possible ^13^C-labeled isotopologues. We quantified NDUFAB1 modifications using the MS2 fragment containing the acyl chain unless noted (Figure S1G). To control for any variation in NDUFAB1-FLAG expression and/or immunoprecipitation efficiency across cell lines, relative NDUFAB1 acylation was calculated by normalizing octanoyl-NDUFAB1 to holo-NDUFAB1. In Mcat-deficient cells, NDUFAB1 acylation is lost, and can be rescued by re-expression of *Mcat* (Figure 1F).

To first test whether malonate was a carbon source for mtFAS, we fed cells uniformly labeled [U-^13^C_3_] malonate and observed strong incorporation into the acyl chain attached to NDUFAB1 in wild type cells (Figure 1G, 1H). Labeling followed a predictable even labeling pattern (M+2, M+4, M+6, M+8) as is typically seen when tracing into cytoplasmic fatty acid synthesis products. Carbon entry was not into the peptide or phosphopantetheinyl (holo) group, as the holo-NDUFAB1 product ion was not labeled (Figure 1I, 1J), and the intact precursors of holo-NDUFAB1 and octanoyl-NDUFAB1 had similar mass isotopologue distributions to their respective product ions (Figure S1H-I). This indicated to us that malonate labels the acyl chain but not the peptide-incorporated amino acids aspartate, serine, or leucine in this time frame (3 hours).

### ACSF3-derived Malonyl-CoA is not required for Mitochondrial Fatty Acid Synthesis

Because we observed label entry from malonate into mtFAS products, we hypothesized that malonate was indeed converted to malonyl-CoA by ACSF3 for consumption by the mtFAS enzyme MCAT. To test this hypothesis, we created CRISPR/cas9-mediated *Acsf3*-knockout clonal cell lines and asked whether mtFAS activity was impaired. Consistent with previous literature^9^ and in contrast to loss of *Mcat*, we saw no decrease in the indirect mtFAS-dependent endpoints protein lipoylation or SDHB stability in *Acsf3*-knockout C2C12 cells (Figure 2A). We also observed no change in the proliferation of *Acsf3*-knockout cells, which similarly contrasts with the phenotypes of other models of mtFAS loss^16^ (Figure 2SA).

**Figure 2.**
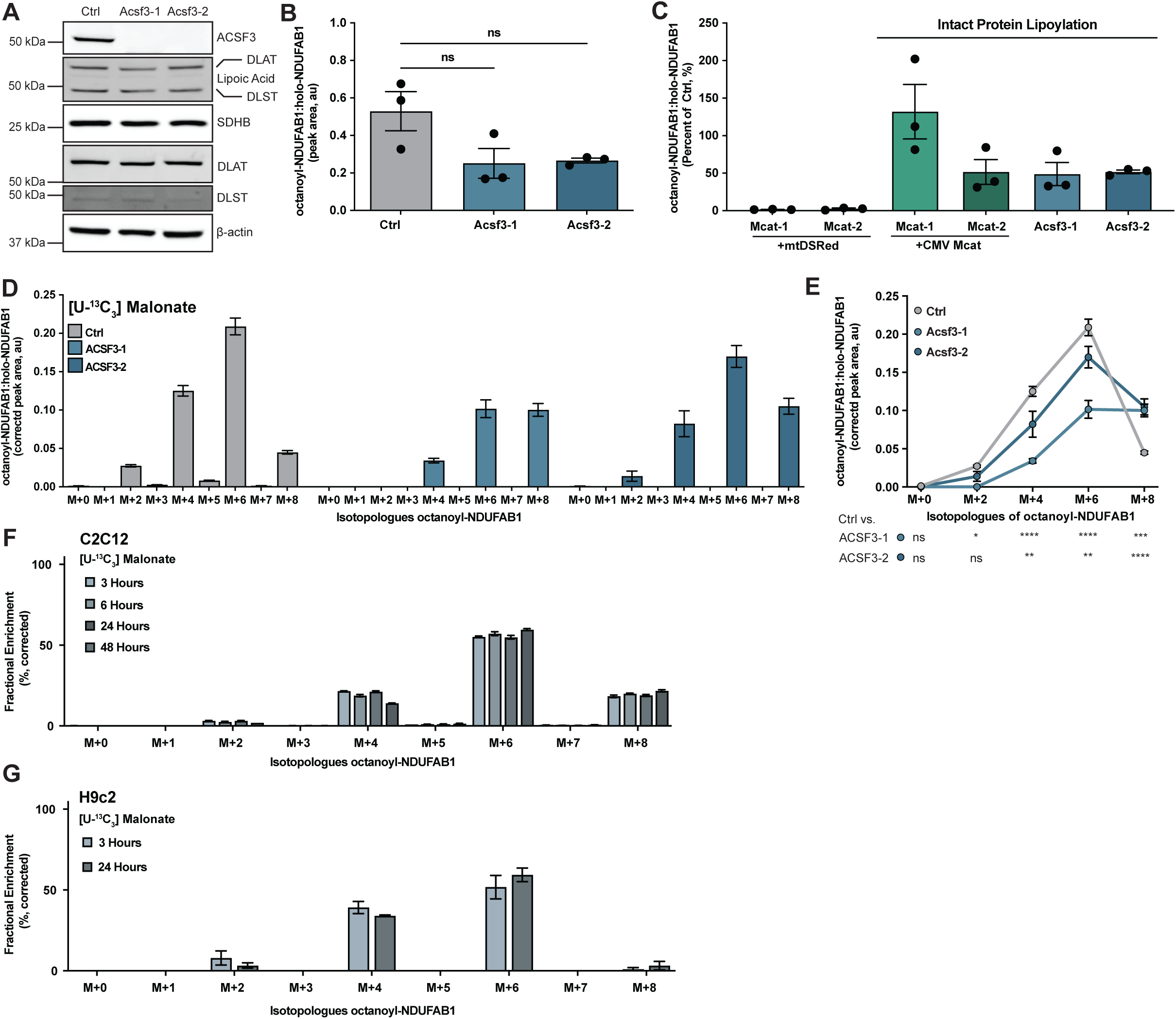
ACSF3-derived Malonyl-CoA is not required for Mitochondrial Fatty Acid Synthesis. (**A**) Whole cell lysates from the indicated cell lines, separated by SDS-PAGE and immunoblotted with the indicated antibodies. (**B**) Relative abundance of octanoyl-NDUFAB1 normalized to holo-NDUFAB1 in the indicated clonal C2C12 cell lines. Statistical analysis performed by one-way ANOVA followed by Tukey’s post-hoc test, (n=3). ns = p>0.05. (**C**) Ratio of octanoyl-NDUFAB1 to holo-NDUFAB1 expressed as a percent of the matched control in the indicated cell line (n=3). (**D**) Mass isotopologue distribution of octanoyl-NDUFAB1 relative to holo-NDUFAB1 in the indicated cell line after culture with uniformly labeled malonate for 24 hours (n=3) (**E**) Overlapping mass isotopologue distribution of even isotopologues from the indicated cell lines. Statistical analysis done by two-way ANOVA with Dunnet’s multiple comparisons test (n=3). ns = p>0.05, * = p<0.05, ** = p<0.01, *** = p<0.001, **** = p<0.0001. (**F**) Mass isotopologue distribution of octanoyl-NDUFAB1 in wild-type C2C12 cells after culture with uniformly labeled malonate for the indicated period of time (n=3), expressed as a percent of total pool (%). (**G**) Mass isotopologue distribution of octanoyl-NDUFAB1 in wild-type H9c2 cells after culture with uniformly labeled malonate for the indicated period of time (n=3), expressed as a percent of total pool (%). Error bars represent +/- 1 SEM.

To directly assess mtFAS activity, we expressed NDUFAB1-FLAG in *Acsf3*-knockout cells (Figure 2SB) and quantified acylation as above (Figure 1E). In *Acsf3*-knockout cells, NDUFAB1 acylation trended lower but was not significantly reduced (p>0.05) compared to controls (Figure 2B). In fact, relative acylation remained comparable to levels observed in our *Mcat* re-expression rescue cell lines, which are also protein lipoylation and ETC assembly competent (Figure 2C). We next asked whether the preservation of NDUFAB1 acylation in the absence of Acsf3 was due to compensation by some other pathway. Feeding labeled malonate (U-^13^C_3_) still resulted in robust labeling of the NDUFAB1-attached acyl chain in the *Acsf3*-knockout cells, at levels only somewhat reduced to that seen in controls (Figure 2D, 2E), indicating that malonate can enter the mtFAS pathway independently of ACSF3.

We also wondered whether malonate was the sole carbon source for mtFAS. To answer this question, we cultured wild type cells in labeled malonate for longer time periods (24 hours for H9c2 or 48 hours for C2C12) and monitored the formation of fully labeled (M+8) octanoyl-NDUFAB1. Remarkably, we observed no changes in the fractional enrichment of octanoyl-NDUFAB1 isotopologues with additional time in label (Figure 2F, 2G). These data indicated that malonate incorporation into octanoyl-NDUFAB1 was already at steady-state after 3 hours. Moreover, we concluded that another carbon source in addition to malonate must support mtFAS activity under these culture conditions since M+4 and M+6 isotopologues are still abundant at steady state (Figure 2F, 2G). Taken together, these results demonstrate the Acsf3-independent incorporation of malonate into mtFAS products and identify the existence of an additional non-malonate carbon source that feeds the pathway.

### Under physiologic conditions cells engage a malonate synthesis pathway to support mtFAS

Cytoplasmic fatty acid synthesis is well known to be fueled by acetyl-CoA, which is converted to malonyl-CoA by ACC1. Because fully labeled (M+8) octanoyl-NDUFAB1 did not accumulate over time in malonate-labeled cells, we asked whether an acetyl-CoA source might provide the remaining carbon in these media conditions by feeding cells uniformly labeled [U-^13^C_6_]-glucose. We found that when malonate is present as in Figure 2E, glucose also labels octanoyl-NDUFAB1 (Figure 3A) with M+2 isotopologues predominating, indicating that glucose provides at least some of the carbon not accounted for by malonate labeling. As with our earlier experiments, this label was observed in the MS2 fragment, indicative of acyl chain labeling.

**Figure 3.**
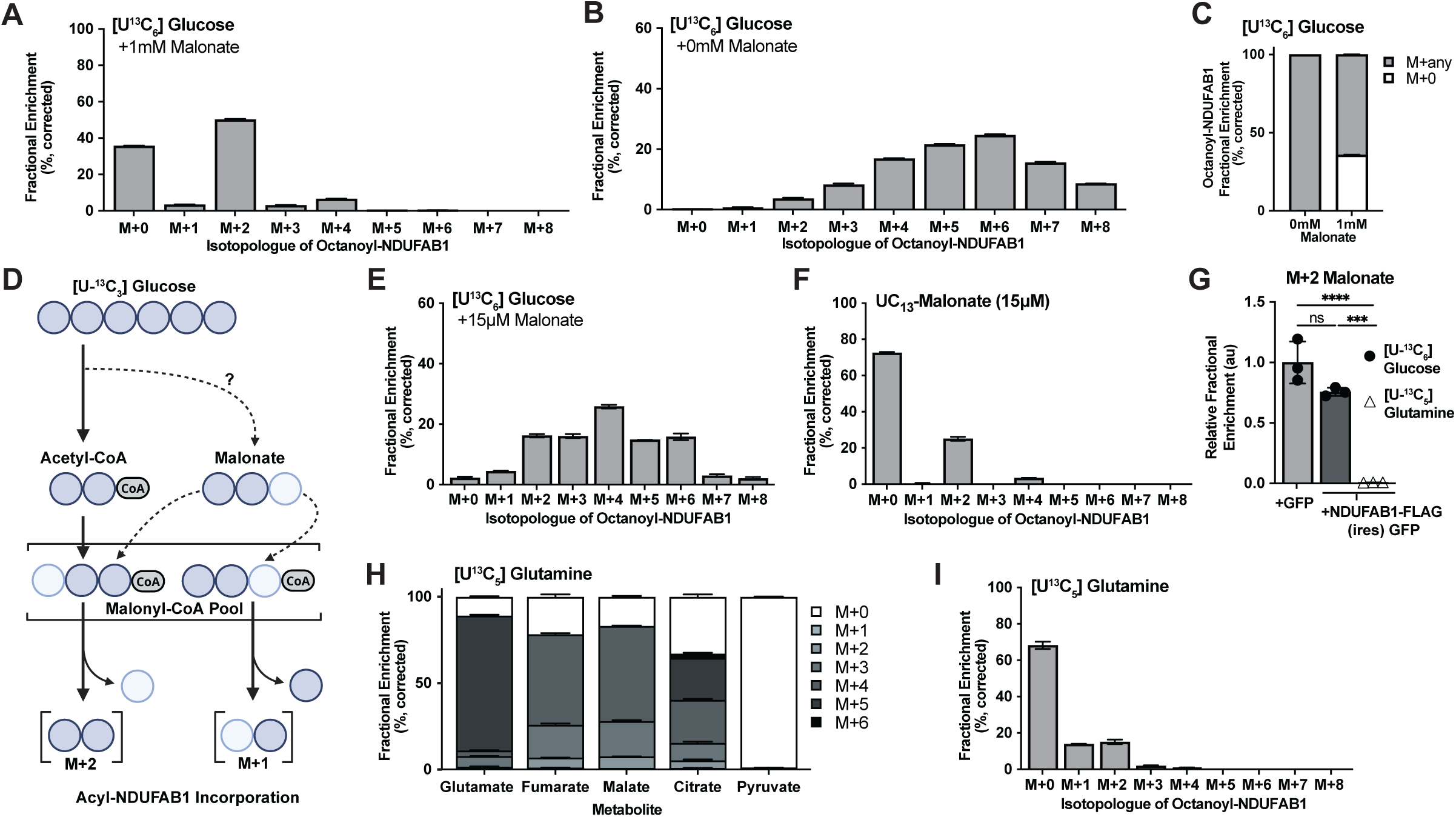
Under physiologic conditions cells engage a malonate synthesis pathway to support mtFAS. **(A**) Mass isotopologue distribution of octanoyl-NDUFAB1 in wild-type C2C12 cells after culture with uniformly labeled glucose for 48 hours in the presence of 1 mM malonate (n=3), expressed as a percent of total pool (%). (**B**) Mass isotopologue distribution of octanoyl-NDUFAB1 in wild-type C2C12 cells after culture with uniformly labeled glucose for 48 hours without no additional malonate (n=3), expressed as a percent of total pool (%). (**C**) Mass isotopologue distributions depicted in (A) and (B) expressed as stacked bars comparing unlabeled octanoyl-NDUFAB1 (M+0) to any other isotopologue of octanoyl-NDUFAB1. (**D**) Diagram depicting metabolic routes to produce observed labeling patterns. Darker circles indicate ^13^C-labeled carbons, lighter circles indicate unlabeled carbons. (**E**) Mass isotopologue distribution of octanoyl-NDUFAB1 in wild-type C2C12 cells after culture with uniformly labeled glucose for 24 hours in the presence of 15 µM malonate (n=3), expressed as a percent of total pool (%). (**F**) Mass isotopologue distribution of octanoyl-NDUFAB1 in wild-type C2C12 cells after culture with 15 µM uniformly labeled malonate for 24 hours (n=3), expressed as a percent of total pool (%). (**G**) Relative fractional enrichment of malonate detected in cells of the indicated genotype from the C2C12 mouse myoblast lineage, labeled with the indicated ^13^C metabolite for 24 hours. Statistical analysis done via one-way ANOVA and Tukey’s post-hoc test (n=3). ns = p>0.05, *** = p<0.001, **** = p<0.0001. (**H**) Fractional enrichment of the indicated metabolite after culture in [U-^13^C_5_]-glutamine for 48 hours. (**I**) Mass isotopologue distribution of octanoyl-NDUFAB1 in wild-type C2C12 cells after culture with uniformly labeled glutamine for 48 hours without the addition of malonate (n=3), expressed as a percent of total pool (%). Error bars represent +/- 1 SEM.

Under standard culture conditions, malonate is not a component of common media formulations (DMEM, RPMI, F12), so we asked whether glucose plays a larger role in mtFAS activity in the absence of exogenous malonate. We found that in wild type C2C12 cells, glucose labeled much more substantially and even reached full labeling (M+8) of the acyl chain (Figure 3B-C) when malonate was not added to the culture media. This was also the case in cultured H9c2 cells grown under the same conditions (Figure S3A, S3B). Surprisingly, the mass isotopologue distribution was significantly affected by the presence or absence of malonate, with the appearance of odd-labeled octanoyl-NDUFAB1 acyl chain isotopologues (M+1, M+3, M+5, M+7) in the absence of malonate (Figure 3B, S3B). This odd labeling pattern indicated to us that the predicted conversion of [U-^13^C_6_]-glucose to [U-^13^C_2_]-acetyl-CoA and then [^13^C_1,2_]-malonyl-CoA could not fully explain the glucose labeling, as this pathway would result in the same even isotopologue pattern as observed for malonate, and classically seen for cytoplasmic fatty acid synthesis (Figure 3D). Rather, this labeling pattern revealed that M+1 incorporations into mtFAS products were taking place. Again, this labeling pattern was not explained by labeling into the digest product’s peptide backbone or phosphopantetheinyl group, as we see no label entry in the holo-NDUFAB1 fragment (Figure S3C). Interestingly, M+1 labeling from glucose also occurred when cells were given exogenous malonate at 15 µM, mimicking physiologic malonate levels and those in human plasma-like medium (HPLM)^17,18^ (Figure 3E, S3D). Labeling exogenous malonate at 15 µM also resulted in label incorporation into the acyl chain, indicating that cells still take up and use exogenous malonate for mtFAS at this lower concentration (Figure 3F, S3E).

We hypothesized that the M+1 labeling of mtFAS products from glucose could be explained if M+2 malonate were being produced, then converted to malonyl-CoA for mtFAS (Figure 3D). Malonate is a symmetrical molecule, therefore CoA can be added to either end of the dicarboxylate. M+2 malonate would therefore give rise to M+1 incorporations if the CoA was added to the side opposite the two carbon-13 atoms on malonate (Figure 3D). However, as discussed above, how malonate might be produced from glucose in mammalian cells is uncertain. Regardless, in support of this idea, wild type cells grown in media lacking exogenous malonate and [U-^13^C_6_]-glucose show appreciable levels of M+2 labeled malonate, and this was not affected by the expression of NDUFAB1-FLAG (Figure 3G).

Glucose readily labels tricarboxylic acid (TCA) cycle intermediates, and one proposed source of cellular malonate is the decarboxylation of the TCA intermediate oxaloacetate (OAA), as described in Fedotcheva et al.^19,20^. To test whether decarboxylation of OAA into malonate was happening in our cells, we labeled TCA intermediates with [U-^13^C_5_]-glutamine. Glutamine efficiently labels TCA cycle metabolites (Figure 3H), however, we observe limited label entry from [U-^13^C_5_]-glutamine into mtFAS products (Figure 3I) and no label entry into malonate (Figure 3G). We therefore concluded that OAA decarboxylation was unlikely to be a major driver of glucose labeling into malonate and downstream malonyl-CoA.

### Endogenous malonate synthesis and mitochondrial oxidative function are dependent on ACC1

To explore what enzymatic sources of malonyl-CoA might be related to mtFAS activity, we analyzed gene co-dependency relationships of known malonyl-CoA-producing enzymes ACC1 (gene name *ACACA*), ACC2 (gene name *ACACB*), and ACSF3 (*ACSF3*) using the DepMap Public 23Q+ Score Chronos dataset^21,22^. Of these three enzymes, only the gene encoding ACC1 was positively correlated with mtFAS genes, including *MCAT*, *MECR*, *OXSM*, and *NDUFAB1* (Figure S4A). We therefore hypothesized that the glucose labeling into mtFAS products was via ACC1, which would produce M+2 malonyl-CoA from labeled acetyl-CoA, followed either by incorporation into the acyl chain or the subsequent loss of CoA to yield M+2 malonate. To test whether ACC1 was required for mtFAS activity, we generated CRISPR/cas9-mediated *Acc1* knockout clonal C2C12 cell lines. As with ACSF3-knockout cells, we observed no significant changes in protein lipoylation or SDHB stability in *Acc1* knockout cells (Figure 4A). However, when we fed these cells [U-^13^C_6_]-glucose, M+2 labeled malonate accumulated in *Acsf3-knockout* cells, but was depleted in *Acc1* knockout cells (Figure 4B). These results demonstrate that ACC1 is required for efficient production of malonate from glucose and suggest that ACSF3 may still play a role in its consumption, even if not significantly contributing to mtFAS products.

**Figure 4.**
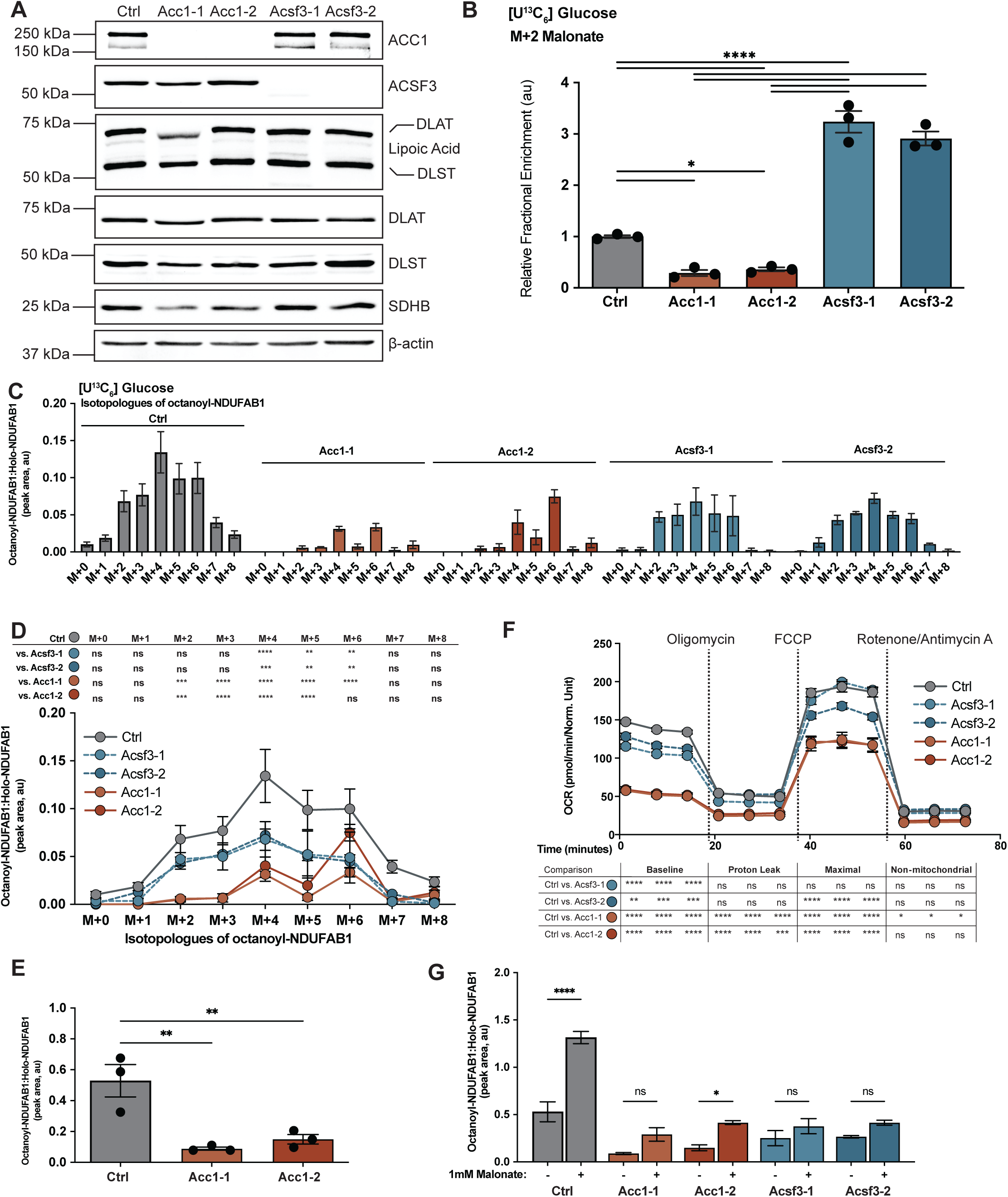
Endogenous malonate synthesis and mitochondrial oxidative function are dependent on ACC1. (**A**) Whole cell lysates separated by SDS-PAGE of the indicated cell line, immunoblotted with the indicated antibody. (**B**) Relative fractional enrichment of M+2 malonate in the indicated cell lines after 24 hours of culture with uniformly labeled glucose for 24 hours (n=3). Expressed as a fraction of M+2 malonate detected in control cell line. Analysis done by one-way ANOVA and Tukey’s post-hoc test. * = p<0.05, **** = p<0.0001. Not significant (p>0.05) comparisons not shown. (**C**) Mass isotopologue distribution of octanoyl-NDUFAB1 in C2C12 cells of the indicated genotype after culture with uniformly labeled glucose for 48 hours without additional malonate (n=3), expressed as octanoyl-NDUFAB1 isotopologue corrected peak area normalized to holo-NDUFAB1. (**D**) Overlapping mass isotopologue distribution of octanoyl-NDUFAB1 isotopologue pool relative to holo-NDUFAB1 from the indicated cell lines. Statistical analysis done by two-way ANOVA with Dunnet’s multiple comparisons test (n=3). ns = p>0.05, * = p<0.05, ** = p<0.01, *** = p<0.001, **** = p<0.0001. (**E**) Relative abundance of octanoyl-NDUFAB1 normalized to holo-NDUFAB1 in the indicated clonal C2C12 cell lines. Statistical analysis performed by one-way ANOVA followed by Tukey’s post-hoc test, (n=3). ** = p<0.01. (**F**) Oxygen consumption rate (OCR) from Seahorse standard mitochondrial stress test, normalized to cell confluency in the indicated C2C12 cell lines. Analysis done by two-way ANOVA with Dunnet’s multiple comparisons test (n=3). ns = p>0.05, * = p<0.05, ** = p<0.01, *** = p<0.001, **** = p<0.0001. (**G**) Relative abundance of octanoyl-NDUFAB1 normalized to holo-NDUFAB1 in the indicated C2C12 cell lines cultured with or without 1mM malonate. Statistical analysis performed by one-way ANOVA followed by Bonferroni’s post-hoc test, (n=3). ns = p>0.05, * = p<0.05, **** = p<0.0001.

We next examined [U-^13^C_6_]-glucose label entry into acyl-NDUFAB1 in our *Acc1* and *Acsf3-*knockout cell lines. We observed octanoyl-NDUFAB1 label incorporation from glucose in both the *Acc1* and *Acsf3-*knockout cells, however *Acc1* knockouts showed notably lower levels of label across all isotopologues (Figure 4C, 4D), as well as significantly reduced total NDUFAB1 acylation (Figure 4E). Interestingly, the labeling pattern in the *Acc1* knockouts also reflects the reduction in glucose-labeled malonate seen in Figure 4B, as M+1 incorporations into acyl-NDUFAB1 are lost. In all cell lines tested, M+1 incorporations were also lost with addition of 1mM exogenous malonate, highlighting how these cells rewire metabolism to support mtFAS depending on malonate availability (Figure S4B). Dual impairment of ACC1 and ACSF3, via siRNA-mediated knockdown of *Acc1* in the *Acsf3*-knockout cells did not diminish protein lipoylation or SDHB stability, further evidence of an additional pathway from malonate to mtFAS (Figure S4C).

Because ACC1 is required for malonate synthesis and downstream mtFAS activity, we wanted to further test if ACC1 was required for mitochondrial oxidative function. We therefore performed a standard mitochondrial stress test on the Seahorse XF, in which mitochondrial respiration is driven by glucose. We found that *Acc1* knockouts displayed reduced oxygen consumption at baseline and at uncoupler (FCCP)-induced maximal respiration (Figure 4F). By contrast, *Acsf3* knockouts had intact oxygen consumption relative to the *Acc1* knockouts (Figure 4F).

We also tested whether adding malonate to cells at either physiologic concentrations (15 μM) or in conditions that reduce malonate synthesis (1 mM), was able to drive mtFAS activity and downstream pathways enough to boost mitochondrial respiratory function. While malonate addition had no effect on baseline oxygen consumption in any of the cell lines, acylation of NDUFAB1 was increased in cells grown in 1 mM malonate, regardless of genotype (Figure 4G, Figure S4D-E). Altogether, these results establish malonate as a critical endogenous intermediate in mitochondrial fatty acid synthesis, demonstrate that cells tune production of endogenous malonate according to the availability of exogenous malonate, and reveal dependencies on ACC1 and an ACSF3-independent pathway to sustain mtFAS function.

## DISCUSSION

Here we identify products of the mtFAS pathway as an important intracellular fate of malonate. This expands the view of malonate beyond its classical role as a toxic complex II inhibitor. Our data show that while cells readily use exogenous malonate when available, under physiologic conditions cells engage a pathway to synthesize malonate from glucose to sustain mtFAS function. These findings establish malonate as a regulated endogenous metabolic intermediate in skeletal and cardiac myoblasts, with important consequences for mtFAS pathway activity and mitochondrial oxidative function.

Our data show that the route from malonate to malonyl-CoA to mtFAS is more complex than previously appreciated, as stable loss of *Acsf3* does not affect the ability of cells to use malonate for mtFAS, directly challenging models that position ACSF3 at the top of the mtFAS pathway. We instead speculate that another mitochondrial enzyme is able to compensate for ACSF3 loss and convert mitochondrial malonate to malonyl-NDUFAB1. While ACSF3 is not required, we observed that the endogenous synthesis of malonate and mtFAS products both depend on ACC1, which converts acetyl-CoA to malonyl-CoA. This is in line with the only other study to show malonate synthesis in a live mammalian system^23^. In that early study, Riley et al. described malonate labeling from exogenous acetate in developing rat brains, in a manner consistent with malonyl-CoA as a metabolic intermediate, but did not observe labeling from citrate or pyruvate. Our data build on these findings, identifying glucose as a physiologically relevant precursor of malonate and establishing the requirement of ACC1 for endogenous malonate synthesis.

In agreement with our data, and further supporting a role for ACC1 upstream of mtFAS, yeast with impaired acetyl-CoA carboxylase function have been shown to display reduced respiratory capacity and a reduction, but not a total loss, of lipoic acid^24^. However, only one report has suggested the possibility that ACC1 may contribute to mtFAS in mammalian cells^25^. Those authors proposed a secondary localization of ACC1 to the mitochondria, enabling the direct use of ACC1-derived malonyl-CoA for mtFAS. While not directly in conflict with a mitochondrial role for ACC1, our data suggest another potential route: if ACC1-derived malonyl-CoA in the cytoplasm is converted to cytoplasmic malonate, then, similar to exogenous malonate, this metabolite could cross the mitochondrial membrane to support mtFAS. This could provide a rationale for why cells would induce a pathway to convert malonyl-CoA to malonate and back again before eventual incorporation into mtFAS products.

Clarifying the compartmental localization of endogenous malonate production will have important implications for considering factors such as substrate availability and additional upstream pathways and metabolites that may feed into endogenous malonate pools. Distinguishing between these models will also require identifying the thioesterase that acts on ACC1-derived malonyl-CoA to produce malonate, as well as the acyl-CoA synthetase that converts malonate back to malonyl-CoA and/or compensates for ACSF3 loss. Defining these enzymes will be essential for understanding the regulation of endogenous malonate synthesis and downstream pathways such as mtFAS.

## DATA AVAILABILITY

All data needed to evaluate the conclusions in the manuscript are present in the manuscript and/or Supplementary Information. Additional data related to this paper may be requested from the authors.

## ACKNOWLEDGEMENTS

This research was funded by the NIH (R35GM151245 to SMN and F30GM154476 to RJW) and the Van Andel Institute (VAI) – Metabolism & Nutrition (MeNu) Program. Stipend support for RJW was provided by the Van Andel Institute Graduate School (VAIGS). We also thank the VAI’s Flow Cytometry Core (RRID:SCR_022685) for their assistance with single cell sorting for generation of clonal H9c2 lines, and GFP-expressing cell lines for NDUFAB1-FLAG pull-down. We thank the Mass Spectrometry Core (RRID:SCR_024903) for their method development of the wide-isolation parallel reaction monitoring mass spectrometry approach.

## AUTHOR CONTRIBUTIONS

Conceptualization (RJW, RDS, SMN)

Formal Analysis (RJW, PRN, MTC)

Funding Acquisition (RJW, SMN)

Investigation (RJW, PRN, MTC, SS, JZL, MD)

Methodology (RJW, PRN, MTC, AEE, RDS)

Resources (AEE, RDS, SMN)

Supervision (RDS, SMN)

Validation (RJW, PRN, AEE, RDS)

Visualization (RJW, PRN)

Writing – original draft (RJW, SMN)

Writing – review & editing (RJW, SMN, AEE, RDS, SMN)

## METHODS

### Cell Culture

C2C12 immortalized mouse skeletal myoblasts (ATCC CRL-1772, verification provided by ATCC) and H9c2 immortalized rat cardiac myoblasts (ATCC CRL-1446, verification provided by ATCC) were grown in DMEM with 4.5 g/L glucose, 4 mM glutamine, and sodium pyruvate (Corning, 10-013-CV) and 10% FBS (Sigma, F0926) at 37°C, 5% CO_2_. For all labeling experiments, cells were cultured in 10% dialyzed serum (Sigma, F0392).

### Generation of knockout cell lines

For each gene targeted (*Mcat*, *Acaca*, and *Acsf3*) sgRNA sequences (Table 1) were designed targeting exon 1 or exon 2 and subcloned into the pLentiCRISPRv2-GFP (addgene #82416) vector^26^. Parental C2C12 or H9c2 cells were transfected with pLentiCRISPRv2_sgRNA targeting luciferase (control) or the gene of interest using JetOptimus reagent (Polyplus, 76299-630). 48 hours after transfection, single GFP+ cells were sorted into 96-well plates to obtain clonal lines. Clones were screened for protein expression; one control clone and two knockout clones of each target were selected for further experiments.

**Table 1.**
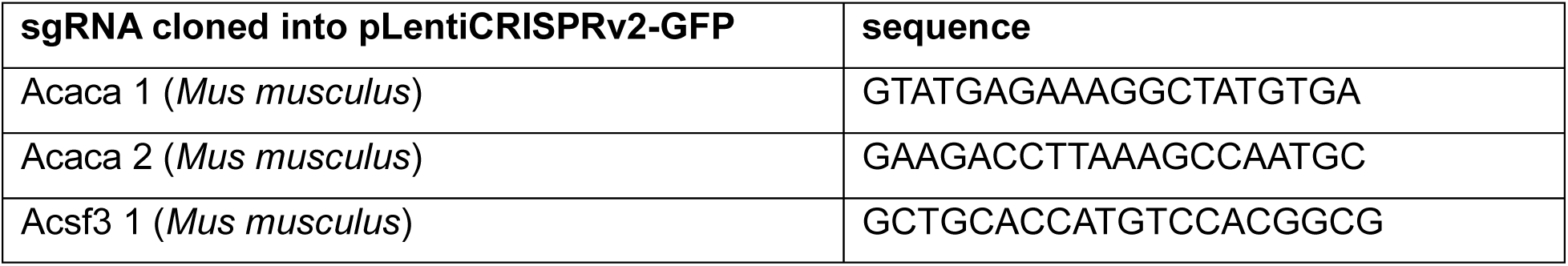
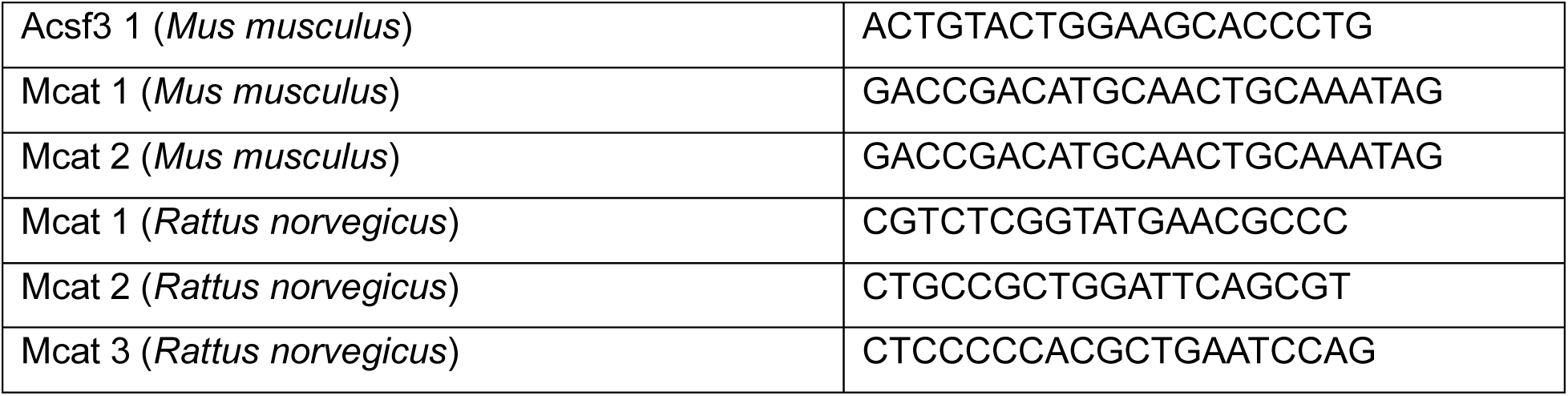

**Table 2.**

| Antibodies | Source and reference number |
| --- | --- |
| ACACA | Cell Signaling 4190 |
| ACSF3 | Santa Cruz sc-398650 |
| ATP5a | Abcam ab14748 |
| β-actin | Proteintech 66009-1-Ig |
| Citrate Synthase | Cell Signaling 14309 |
| DLAT | Abcam ab172617 |
| DLST | Cell Signaling 5556 |
| DYKDDDK (FLAG) | Invitrogen MA1-91878 |
| GFP | Abcam ab290 |
| GRIM19 | Abcam ab110240 |
| Lipoic acid | Millipore 437695 |
| MCAT | Santa Cruz sc-390858 |
| MTCO1 | Abcam ab110270 |
| NDUFA9 | Abcam ab14713 |
| NDUFAB1 | Abcam ab96230 |
| SDHA | Abcam ab14715 |
| SDHB | Abcam ab14714 |
| UQCRFS1 | Abcam ab14746 |

### Retroviral-Mediated Gene Expression

For rescue experiments, control and knockout lines were infected with control retrovirus (pQXCIP mtDSRed) or pMMLV retrovirus vectors conferring puromycin resistance (purchased from VectorBuilder; Chicago, IL, USA), harboring the indicated species-specific gene under a CMV promoter. Cells were then selected using 2 μg/mL puromycin. For NDUFAB1-FLAG expression, control and knockout lines were infected with control retrovirus encoding GFP alone, or NDUFAB1-FLAG followed by an IRES-GFP cassette. Cells were then selected for by flow-sorting for GFP expression levels.

### Crude mitochondrial isolation

Cells were harvested and processed as previously described^16^. Briefly, cells were harvested by trypsinization, washed once with cold, sterile PBS (Gibco, 10010-023), then stored at -80°C. At the time of mitochondrial isolation, cell pellets were thawed and resuspended in 1 mL CP-1 buffer (100 mM KCl, 50 mM Tris-HCL, 2 mM EGTA, pH 7.4) supplemented with mammalian protease inhibitor cocktail (mPIC, Millipore Sigma P8340) and mechanically lysed by 10 passes through a 1 mL insulin syringe with 26-guage needle and centrifuged at 700 x g to pellet unlysed cells and debris. Supernatant was transferred to a new microcentrifuge tube and centrifuged at 10,000 x g to pellet the crude mitochondrial fraction. Mitochondrial pellets were resuspended in a small volume of either RIPA buffer (10 mM Tris-HCl, pH 8.0, 1 mM EDTA, 0.5 mM EGTA, 1% Triton X-100, 0.1% Sodium Deoxycholate, 0.1% SDS, 140 mM NaCl) supplemented with mPIC, or CP-1 buffer, equal to approximately twice the pellet volume. Resuspended pellets were then utilized for applications described below.

### SDS-PAGE and immunoblotting

Cultured cells were scraped from tissue culture plates directly into RIPA buffer supplemented with mPIC and Pierce Universal Nuclease, incubated at 4°C with constant agitation on a microcentrifuge tube orbital shaker, then centrifuged at 17,000 x g for 20 minutes at 4°C to remove insoluble material. Supernatant was then transferred to a new microcentrifuge tube to be saved as whole cell lysate (WCL). For denoted protein analysis, whole cell lysates and/or crude mitochondrial fractions were first normalized for total protein content via Pierce BCA Protein assay (Thermo Fisher Scientifc 23225). Samples were resolved by SDS-PAGE and transferred to nitrocellulose membranes. Immunoblotting was performed using the indicated primary antibodies (Table 3) with visualization and analysis of protein signal using Rabbit and Mouse IgG Antibody DyLight 680, or 800, conjugated secondary antibodies and Bio-Rad ChemiDoc Imaging System.

**Table 3.**
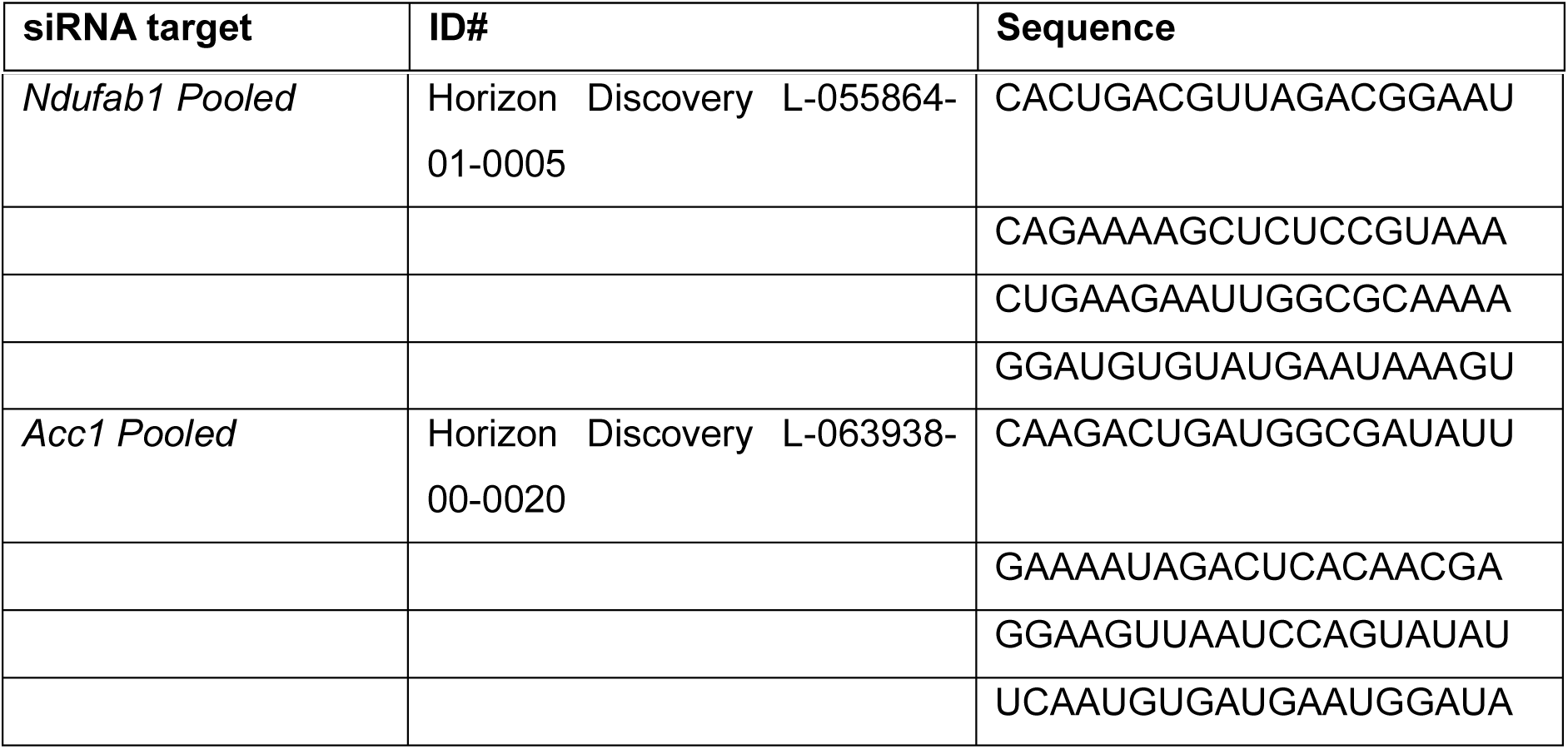

### Blue Native-PAGE

Crude mitochondrial fractions were prepared and normalized for total protein content as previously described. 100 μg of mitochondria were pelleted at 10,000 x g for 10 minutes at 4°C. Supernatant was removed and pellets were resuspended in 60 μL of 1x pink lysis buffer (Invitrogen BN2003). Digitonin (GoldBio D-180-2.5) was added to samples at a final concentration of 1% mass/volume and samples were incubated on ice for 15 min. Insoluble material was pelleted by centrifugation at 17,000 x g for 20 minutes and supernatant was transferred to new microcentrifuge tubes. 6 μL of NativePAGE sample buffer (Invitrogen BN2004) was added to samples and 10 μL of sample (∼15 μg of crude mitochondrial fraction) was loaded per lane on pre-cast 3-12%, Bis-Tris NativePAGE gels (Invitrogen BN1001BOX) with NativePAGE anode buffer (Invitrogen 2001) and dark blue cathode buffer (Invitrogen BN2002) at 150 V for 30-40 minutes then switched to light blue cathode buffer and run at 20 V overnight. Gels were then transferred to PVDF membranes at 100 V for 2 hours and blotted with the indicated primary antibodies (Table 3). Secondary anti-mouse HRP antibody and SuperSignal West Femto Maximum Sensitivity Substrate (Thermo Fisher Scientific 34096) was used to visualize bands with a Bio-Rad ChemiDoc Imaging System.

### Extracellular flux Analysis

Oxygen consumption assays were performed on a Seahorse XFe96 Analyzer (Agilent). Assays were performed in seahorse media (Agilent 103680) with 25 mM glucose, 2 mM glutamine, and 1 mM sodium pyruvate. Standard mitochondrial stress tests were conducted using 1 mM oligomycin, 3 mM FCCP, and 0.5 mM Rotenone + 0.5 mM Antimycin A. Measurements were taken over 3 minutes, with three measurements per phase. Data were normalized to crystal violet staining for approximation of cell number or by measurement of well % confluency using an IncuCyte® chamber. Results were analyzed in WAVE software and the Agilent Seahorse Analytics browser-based application.

### Carbon Tracing into Metabolites

Cell lines were plated in triplicate in 6-well plates and allowed to adhere overnight, at which point the media was changed to unlabeled tracing media (Gibco DMEM A14430-01 with 1 mM sodium pyruvate, 5 mM glucose, and 4 mM glutamine). Wells were changed to labeled media at the indicated time points prior to harvest, Gibco DMEM A14430-01 with 1 mM sodium pyruvate, 5 mM glucose or 5 mM [U-^13^C_6_]-glucose (Cambridge Isotope Laboratories CLM-1396), and 4 mM glutamine or 4 mM [U-^13^C_5_]-glutamine (Cambridge Isotope Laboratories CLM-1822). Three wells of the control cell line were left unlabeled. At time of harvest, plates were washed with normal saline and frozen at -80°C. Plates were extracted for metabolites using 40:20:20 methanol:acetonitrile:water scaled to the number of cells. Lysates were sonicated and incubated on ice for 60 minutes, then insoluble material was pelleted. 1000 µL of supernatant was dried using a SpeedVac concentrator.

Dried metabolomics extracts were resuspended in 100% water containing 25 µg/mL D5-Glutamate (DLM-556, Cambridge). 2 µL of resuspended samples were injected on column. Data were collected on a Vanquish liquid chromatography system coupled to an Orbitrap Exploris 240 (Thermo Fisher Scientific) using a heated electrospray ionization (H-ESI) source in ESI negative mode. All samples were run through a 24-minute reversed-phase chromatography ZORBAX extend-C18 column (1.8 μm, 2.1 mm × 150 mm, 759700-902, Agilent, California, USA) combined with a guard column (1.8 μm, 2.1 mm × 5 mm, 821725-907, Agilent). Full scan data were collected with a scan range of 70-800 m/z at a mass resolution of 240,000. Fragmentation data was collected using a data-dependent MS2 (ddMS2) acquisition method with MS1 mass resolution at 120,000, MS2 mass resolution at 15,000, and HCD collision energy fixed at 30%. Data analysis was conducted in Skyline (version 23.1.0.268), consisting of peak picking and integration, using an in-house curated compound database of accurate mass MS1 and retention time derived from analytical standards and/or MS2 spectral matches for the specified chromatography method. Natural isotope abundance correction was done using the FluxFix Isotopologue Analysis Tool*14* (version 0.1) web-based application^27^ or IsoCorrectoR^28^.

### Relative quantification of NDUFAB1 species

Method was adapted from^13,14^ as in Norden et al^15^. Standards for NDUFAB1 modifications were synthesized as in Nam et al.^14^ and digested with AspN (Sigma, 11420488001) overnight at 37°C in a ratio of 1:20 (w/w) enzyme to sample, buffered to a pH of 7.6 with MOPS, then quenched by addition of methanol to 50% (v/v). Digested standards were pooled for detection by LC-MS.

Cells of the indicated genotype expressing NDUFAB1-FLAG were seeded in three 150 mm tissue culture plates each to be 60-70% confluent at the time of harvest. Cells were harvested by washing with ice-cold PBS then freezing at -80 degrees Celsius. After being frozen for 16 hours or more, cells were scraped into lysis buffer (1% triton, 10 mM HEPES, 0.3 mM EDTA, 120 mM NaCl, and mPIC) and incubated at 4 degrees Celsius for 20 minutes. Samples were then spun to pellet debris and cleared supernatants were incubated with anti-FLAG magnetic beads (Sigma, M8823). Bound proteins were eluted from beads using Flag peptide (Sigma, F4799) in elution buffer (10mM HEPES, 50mM NaCl), spin-filtered (Sigma, CLS8163), and quantified via Bradford assay (BioRad, 5000205). Eluted protein from each sample were digested with AspN as above. Samples were dried, resuspended in 50% methanol:50% water at a concentration of 0.05 ng/uL, and 10uL of sample was injected into a Vanquish (Thermo Scientific) liquid chromatography system coupled to an Orbitrap ID-X Tribrid mass spectrometer (Thermo Scientific). A reversed-phase C18 column (Agilent ZORBAX RRHS Extend-C18 Column, 759700-902) was used to separate samples using acidic (0.1% formic acid) solvents. The first 1.5 minutes of column eluants were diverted to waste due to the presence of MOPS buffer in this time frame. Mass spectrometer was set to collect data using a parallel-reaction monitoring (PRM) approach to isolate a range or ions of interest over retention times that matched those observed in the standards. The PRM isolation window was centered on a mass to charge ratio of M+ ½n and spanned the entire range of isotopologues possible for a given NDUFAB1 digest product (M+0 to M+n, where n equals the number of carbons in the MS1 precursor ion). Over the same retention time window, a given PRM window was set to a collision energy of either 1 or 30 (**Figure S1G**). Data analysis was conducted in Skyline (version 23.1.0.268).

### Carbon tracing into NDUFAB1 species

As above, cells of the indicated genotype expressing NDUFAB1-FLAG were seeded in three 150 mm tissue culture plates to be 60-70% confluent at the time of harvest. Plates were changed to labeled media at the indicated time points prior to harvest; Gibco DMEM A14430-01 with 1 mM sodium pyruvate, 5 mM glucose or 5 mM [U-^13^C_6_]-glucose (Cambridge Isotope Laboratories CLM-1396), and 4 mM glutamine or 4 mM [U-^13^C_5_]-glutamine (Cambridge Isotope Laboratories CLM-1822). Malonic acid was added as indicated, either 15 µM or 1mM, malonic acid or [U^13^C_3_]-malonic acid (Cambridge Isotope Laboratories, CLM-6123). Cells were harvested by washing with ice-cold PBS followed by freezing at -80 degrees Celsius. NDUFAB1-FLAG immunoprecipitation, digestion, and analysis by mass spec was as above for relative quantification of NDUFAB1 species. Natural isotope abundance correction was done either using IsoCor (v2.2.3)^29^ or (if indicated) the FluxFix Isotopologue Analysis Tool*14* (version 0.1) web-based application^27^.

### siRNA knockdown

To knockdown *Acc1* or *Ndufab1*, siRNAs were acquired as indicated in (**Table 3**). siRNAs were packaged with Lipofectamine RNAiMAX, transfected into C2C12, and cells were cultured for 72 hours post-transfection before harvest. ON-TARGETplus Non-targeting Control Pool (D-001810-10-05) was used as a negative control.

### Statistical analysis

Statistical analysis was performed using GraphPad Prism 10. Data were analyzed by one-way ANOVA followed by Dunnett’s multiple comparison test (when compared to only control) or Bonfferoni’s or Tukey’s multiple comparison test (when comparing all groups). A p-value of <0.05 was considered to be statistically significant.

## Supplementary Figure Legends

**Figure S1.**
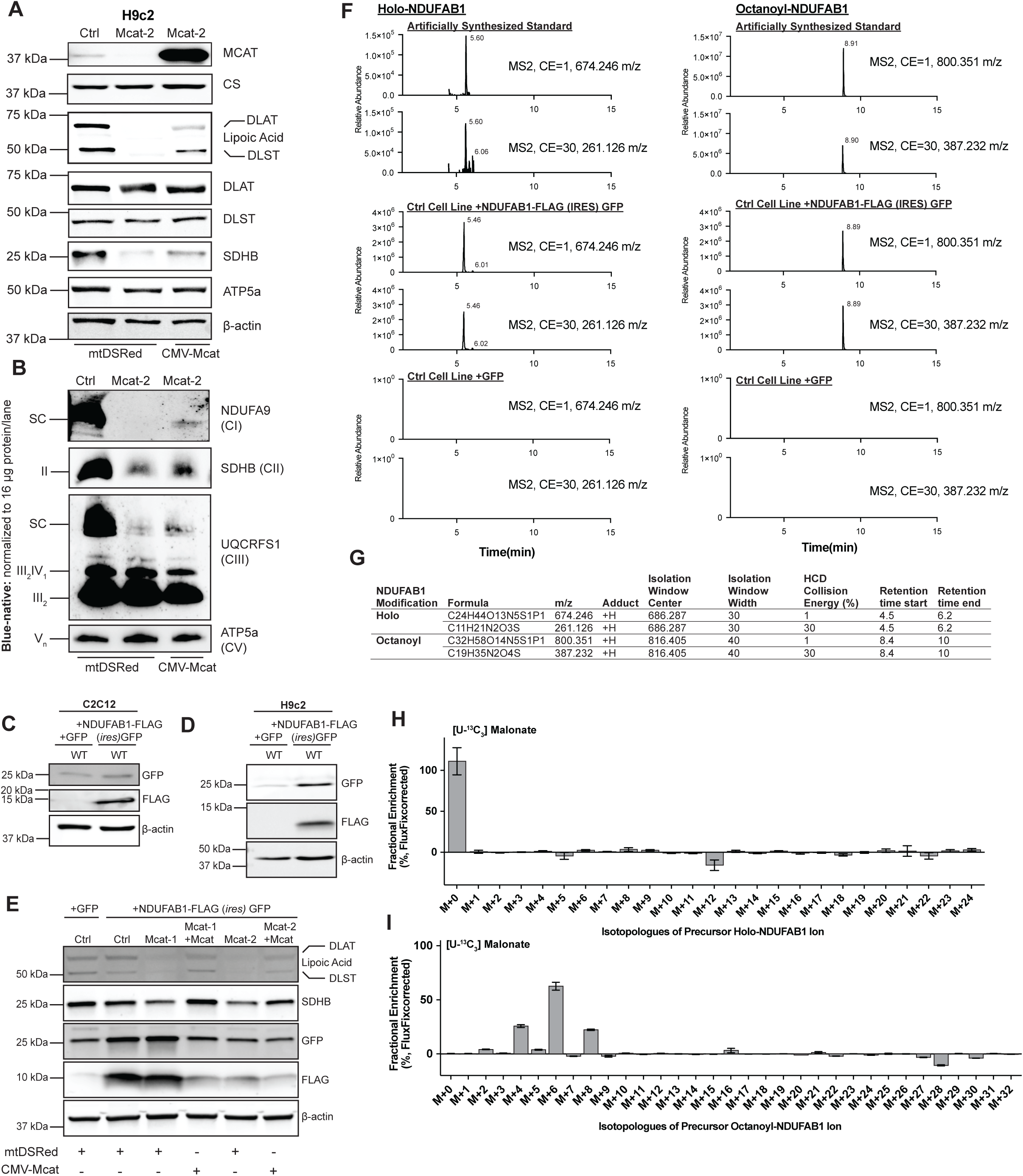
Malonate Drives Mitochondrial Fatty Acid Synthesis. (A) Whole cell lysates or mitochondrial enriched lysates (MCAT, CS) in clonal H9c2 cell lines of the indicated genotypes. (B) Blue-native PAGE of crude isolated mitochondrial lysates from the indicated H9c2 cell lines expressing mtDSRed or Mcat, immunoblotted for the indicated ETC complex subunits. Data are representative of 3 biological replicates. (C-E) Whole cell lysates from (C) wild-type C2C12 cells or (D) wild-type H9c2 cells or (E) C2C12 cells of the indicated genotype expressing mtDSRed or Mcat, expressing GFP or NDUFAB1-FLAG(ires)GFP blotted with the indicated antibodies. (F) Extracted ion chromatograms of ions corresponding to holo-NDUFAB1 or octanoyl-NDUFAB1 from the indicated sample and HCD collision energy (%). (G) Table of quantified ions for the indicated NDUFAB1 modification after AspN digest and the corresponding approach by mass spectrometry for detection in a 15-minute C18 reversed phase liquid chromatography method. (H) Mass isotopologue distribution of precursor ion for holo-NDUFAB1 detected in control cell line (n=3) after culture in uniformly labeled ^13^C malonate for 3 hours. (I) Mass isotopologue distribution of precursor ion for octanoyl-NDUFAB1 detected in a control cell line (n=3) after culture in uniformly labeled ^13^C malonate for 3 hours, corrected with unlabeled sample. Error bars represent +/- 1 SEM.

**Figure S2.**
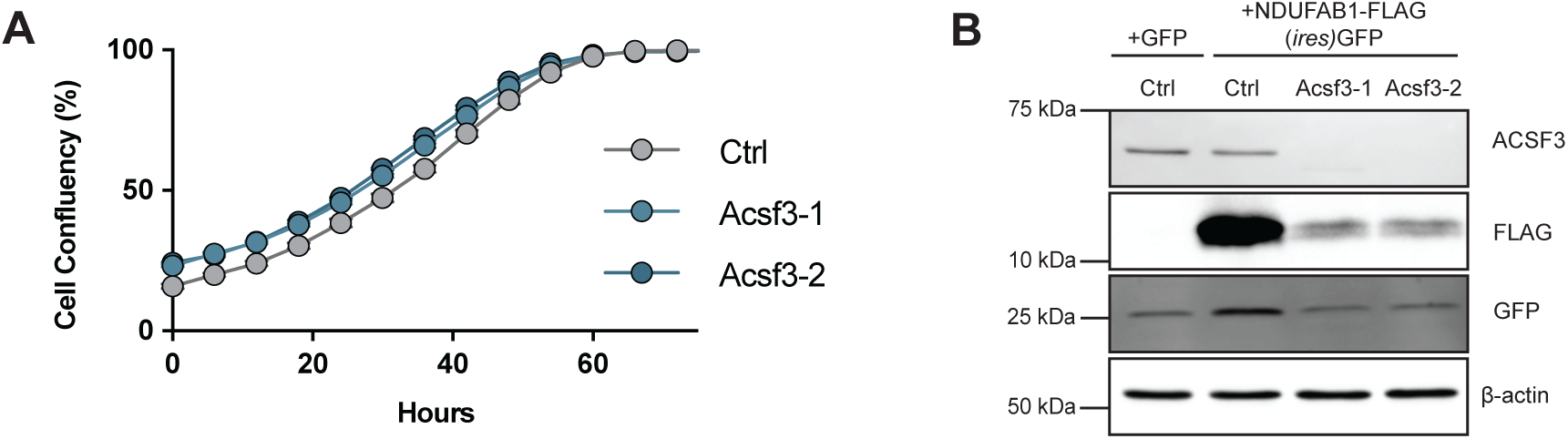
Malonate Drives Mitochondrial Fatty Acid Synthesis. (A) Cells of the indicated genotype seeded in 6-well plates and incubated for the indicated time and assessed for confluency by Incucyte image analysis (n=3). Representative of three biological replicates. (B) Whole cell lysates separated by SDS-PAGE of the indicated cell line, immunoblotted with the indicated antibodies. Representative of three biological replicates. Error bars represent +/- 1 SEM.

**Figure S3.**
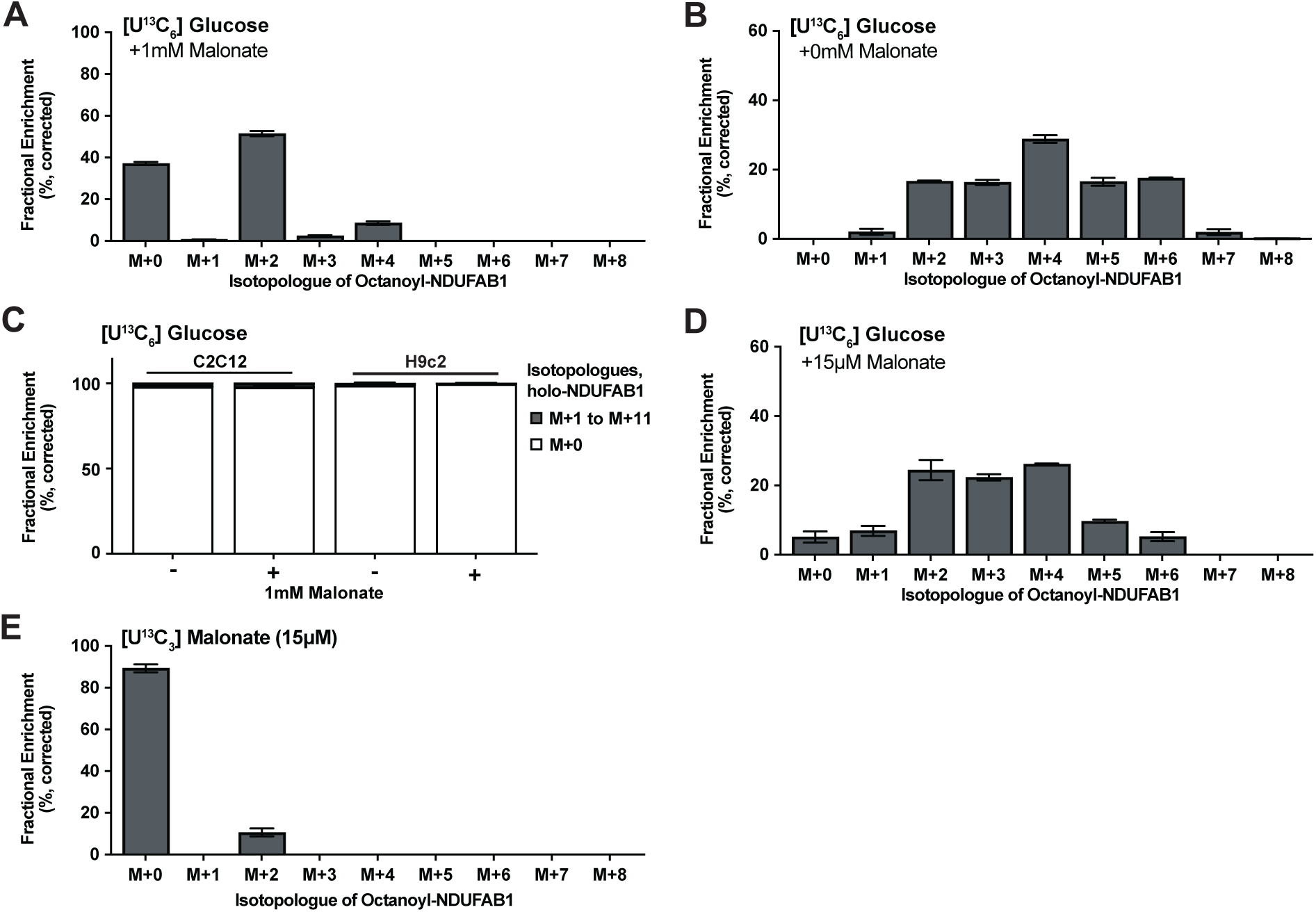
Under physiologic conditions cells engage malonate synthesis to support mtFAS. (A) Mass isotopologue distribution of octanoyl-NDUFAB1 in wild-type H9c2 cells after culture with uniformly labeled glucose for 48 hours in the presence of 1 mM malonate (n=3), expressed as a percent of total pool (%). (B) Mass isotopologue distribution of octanoyl-NDUFAB1 in wild-type H9c2 cells after culture with uniformly labeled glucose for 48 hours without additional malonate (n=3), expressed as a percent of total pool (%). (C) Fractional enrichment (%) of all isotopologues of holo-NDUFAB1 in cells of the indicated lineage grown with or without 1 mM malonate and labeled with [U-^13^C_6_]-glucose for 48 hours (n=3). (D) Mass isotopologue distribution of octanoyl-NDUFAB1 in wild-type H9c2 cells after culture with uniformly labeled glucose for 24 hours in the presence of 15 µM malonate (n=3), expressed as a percent of total pool (%). (E) Mass isotopologue distribution of octanoyl-NDUFAB1 in wild-type H9c2 cells after culture with 15 µM uniformly labeled malonate for 24 hours (n=3), expressed as a percent of total pool (%). Error bars represent +/- 1 SEM.

**Figure S4.**
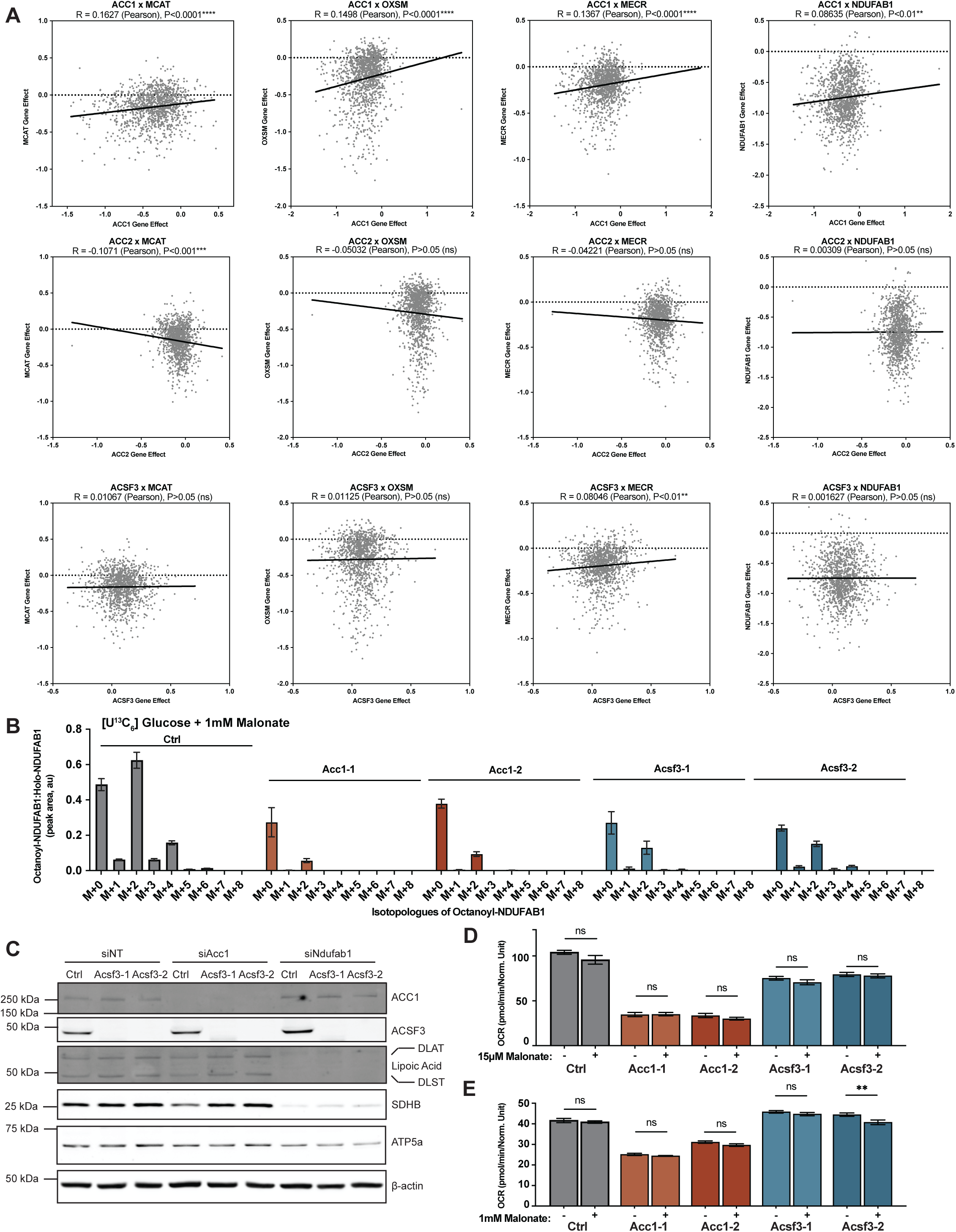
Endogenous malonate synthesis and mitochondrial oxidative function are dependent on ACC1. **(A)** Codependency plots of gene essentiality scores comparing malonyl-CoA producing enzymes ACC1, ACC2 and ACSF3 with genes of the mtFAS pathway: MCAT, OXSM, MECR, and NDUFAB1 from DepMap CRISPR Screens^21,22^. (B) Mass isotopologue distribution of octanoyl-NDUFAB1 in C2C12 cells of the indicated genotype after culture with uniformly labeled glucose for 48 hours with the addition of 1mM malonate (n=3), expressed as a ratio of octanoyl-NDUFAB1 isotopologue corrected peak area to holo-NDUFAB1 peak area. (C) Whole cell lysates of the indicated cell line treated with the indicated gene-targeting siRNA pool for 72 hours. Separated by SDS-PAGE and blotted with the indicated antibodies. (D) Seahorse extracellular flux assay measurement of oxygen consumption rate (OCR) in the indicated cell line of the C2C12 lineage with or without 15 µM malonate treatment overnight (approximately 18 hours), representative of three biological replicates. (E) Seahorse extracellular flux assay measurement of oxygen consumption rate (OCR) in the indicated cell line of the C2C12 lineage with or without 1 mM malonate treatment overnight (approximately 18 hours), representative of three biological replicates. Error bars represent +/- 1 SEM.

